# Gap Junction Protein Connexin43 (Cx43) Promotes Glioma Invasion via the Annexin A2/MMP3 Signaling Axis

**DOI:** 10.64898/2026.09.14.751039

**Authors:** Corbin Glufka, Emmanuel Ojefua, Leonard J. Foster, Wun Chey Sin, Christian C. Naus, Vincent C. Chen

## Abstract

Glioblastoma is among the most aggressive primary brain tumors, with the invasion of surrounding tissue driven by coordinated remodeling of the extracellular matrix (ECM) by proteolytic enzymes, including the matrix metalloproteinase (MMP) family. The gap junction protein connexin43 (Cx43) has emerged as an unexpected promoter of this invasive phenotype. However, the downstream molecular mechanisms linking Cx43 to ECM proteolysis remain poorly understood. We previously demonstrated selective upregulation of MMP3 within the secretome of Cx43-overexpressing glioma cells, implicating Cx43 in the regulation of ECM protease activity. To identify the molecular intermediaries responsible, we carried out a proteomic survey of gap junction (GJ)-associated signaling networks. Amongst these, we identified a Cx43 protein-protein interaction involving annexin A2 (ANXA2), a known regulator of ECM proteases. SDS-PAGE/Western blot and confocal imaging demonstrated that elevated Cx43 expression drives selective recruitment of ANXA2 to gap junction membranes, forming a junctional signaling platform rather than increasing total ANXA2 abundance. This recruitment coincided with co-localization of S100A10, suggesting assembly of an intact ANXA2/S100A10 complex at gap junctions. Using ANXA2-mimetic peptides targeting the S100A10-binding domain, we demonstrate that disruption of the ANXA2/S100A10 complex leads to a significant reduction in MMP3 activity. Furthermore, in 3D spheroids grown in ECM-replicating gels, ANXA2 peptide treatment significantly reduced the invasive potential of C6-13 (Cx43-overexpressing) cells. Together, these data define a novel Cx43→ANXA2→S100A10→MMP3 signaling axis, advancing our understanding of ECM remodeling in glioblastoma and establishing the ANXA2/S100A10 complex as a promising therapeutic target for disrupting Cx43-driven glioblastoma progression.

## Introduction

Gliomas are the most common tumors of the central nervous system (CNS) accounting for ∼80% of malignancies (Holland 2000). Patients diagnosed with glioblastoma (GBM), the highest-grade glioma (grade 4), continue to face poor prognoses with a median overall survival of 12-15 months (Demuth and Berens 2004; Ostrom et al. 2014; Furnari et al. 2007; Weller et al. 2015). Believed to arise from glial precursors or dedifferentiated cells (Parkin et al. 2001; Chen et al. 2017), GBM tumors are characterized by angiogenesis, proliferation, resistance to cell death, and a high propensity to invade surrounding tissue of the CNS. Although a significant portion of GBM tumors can be removed with surgery, diffusely invading cells routinely evade resection, leading to tumor recurrence at secondary sites (Iwadate 2016; Cooper et al. 2012).

The migratory and invasive capacity of GBM is critically dependent on the ability of tumor cells to degrade and remodel the extracellular matrix (ECM), a three-dimensional noncellular network of macromolecules that forms a natural barrier to cell migration (Aftab et al. 2019; Bellail et al. 2004; Giese and Westphal et al. 1996; Mohiuddin and Wakimoto 2021; Nakada et al. 2007). The most significant class of proteases associated with ECM degradation are matrix metalloproteinase (MMP) family, a diverse group of 23 zinc-dependent endopeptidase whose expression is frequently dysregulated in GBM, leading to abnormal ECM degradation and remodeling (Cawston and Young 2010; Cabral-Pacheco et al. 2020; Kapoor et al. 2016). Historically, MMPs were thought to only participate at the beginning of metastasis, however, it is now known that MMPs are involved at all steps of tumor progression, playing a role in the modification of signaling pathways, regulation of immune response and tumor growth (Cabral-Pacheco et al. 2020). Despite this, the upstream signals that drive aberrant MMP activity in GBM remain incompletely understood, and identifying these regulators represents a key step for developing anti-invasive therapies.

One candidate regulator of aberrant MMP activity in GBM is the gap junction (GJs) forming protein Connexin 43 (Cx43) (Liang et al. 2020; Plotkin and Bellido 2013). Canonically, neighboring Cx43s fuse to form densely packed ion channel arrays (plaques) to mediate gap junctional intercellular communication (GJIC) scaffolding (Goodenough and Paul 2009; Naus and Laird 2010; Qin et al. 2016; Zhao et al. 2015). In addition to its channel-forming role, the cytoplasmic carboxy-terminal tail (CT) has been linked to the regulation of non-channel functions such as protein-protein interactions and recruitment of cytoskeletal (Dong et al. 2017; Duffy et al. 2002; Hervé et al. 2007; Laird 2010; Sharrow et al. 2008; Sorgen et al. 2018; Thévenin et al. 2013). Although Cx43 was initially characterized as a tumor suppressor based on early studies showing that its upregulation slowed glioma growth (Fu et al. 2004; Huang et al. 1998; Zhu et al. 1991), subsequent work has demonstrated that Cx43 has a contrasting role as a promoter of glioma malignancy by increasing migration and invasion and providing therapeutic resistance against the front-line chemotherapeutic agent Temozolomide (TMZ) (Aftab et al. 2019; Gielen et al. 2013; Kameritsch et al. 2012; Munoz et al. 2014; Naus et al. 2016; Pridham et al. 2022). Our lab previously demonstrated Cx43-mediated invasion is associated with the activation of the ECM protease MMP-3 (Aftab et al. 2019). Although the pro-invasive function of Cx43 has been localized to the cytoplasmic CT domain (Cx43CT), the downstream molecular pathways by which Cx43CT coordinates invasion-promoting signaling and its links to MMP3 activation remain uncharacterized (Bates et al. 2007; Behrens et al. 2010; Cina et al. 2009; Machtaler et al. 2011; Yilmaz and Christofori 2009).

In this study, we carried out a multi-proteomic approach to characterize the gap junction signaling nexus in a Cx43-overexpressing rat glioma model (C6 and its clone C6-13) with the goal of identifying the molecular intermediaries linking Cx43 to MMP3-driven ECM remodeling (Zhu et al. 1991). This analysis identified annexin A2 (ANXA2) as a candidate Cx43-interacting protein and regulator of the ECM. We provide the first functional evidence that a Cx43→ANXA2→S100A10→MMP3 signaling axis promotes glioma invasion and demonstrate that targeted disruption of the ANXA2/S100A10 complex represents a promising therapeutic strategy for limiting Cx43-driven glioblastoma progression.

## Material and Methods

### Cells and Cell Culture

Mammalian (Rat) C6 and C6-13 (C6-Cx43) glioma cells were cultured in 4.5mg/ml Dulbecco’s modified Eagle medium (DMEM) supplemented with 10% fetal bovine serum (FBS) and 100 U/mL penicillin-streptomycin (PS) antibiotics at 37°C with 4% CO2. The C6-13 cell line is a stable clonal variant derived from parental C6 cells that is engineered to constitutively overexpress Cx43. Cells were passed at 80-90% confluency or every 3 days by aspirating culture media, washing with PBS and then incubated in trypsin-EDTA for 5 min at 37°C. The trypsinized cell suspension was centrifuged for 5 min at 1000 x g, the supernatant aspirated, and the resulting cell pellet resuspended in fresh culture media.

### Gap Junction-enriched Plasma Membrane Preparation

Confluent C6 and C6-13 cultures were washed with ice-cold PBS and scraped into gap junction buffer (5 mM Tris-HCl, 5 mM EDTA, 5 mM EGTA, pH 8.0, 1 mM PMSF). Cells were homogenized using a chilled dounce homogenizer (40–50 strokes), briefly sonicated (10–15 s), and centrifuged at 1000 × g for 10 min at 4°C. Supernatants were collected and centrifuged at 30,000 × g for 1 h at 4°C. Pellets containing GJ-enriched membranes were rinsed with cold buffer and air-dried.

### Peptide Sample Preparation and LC-MS/MS

Gap junction-enriched protein pellet were solubilized in digestion buffer (1% deoxycholate, 50 mM NH₄HCO₃), normalized by BCA assay, heated (99°C, 5–10 min), reduced with DTT (1:50, DTT:protein, 20 min, 37°C), and digested overnight with trypsin (1:50, enzyme: protein). Peptides were acidified, clarified, desalted using STAGE-Tips, and eluted with 80% acetonitrile/0.5% acetic acid. Samples were dried and resuspended in 0.5% acetic acid. Peptides were analyzed on an Agilent 1200 HPLC coupled to an Agilent 6530 QTOF mass spectrometer operating in data-dependent acquisition mode (1 MS scan followed by 5 MS/MS scans).

### Protein Identification and GJ Network Enrichment Analysis

Raw MS files were converted to mgf format using MSConvert and searched using Mascot (v2.6) or directly analyzed using MaxQuant. Mascot parameters included: trypsin/P specificity, UniProt rat database, precursor tolerance 20 ppm, fragment tolerance 0.6 Da, carbamidomethyl (C) as a fixed modification, and methionine oxidation as a variable modification. Data sets encompassing gap junction-enriched membranes identified 1737 proteins. This bioinformatics analysis was further enabled using a previous Cx43-protein interaction study (Chen et al. 2012). This data was extracted from the BioGRID database. Interactome identifiers used here maybe obtained from the original paper (IPI) or downloaded (Uniprot) from BioGrid (thebiogrid.org, publication search ID: 23106098). Statistical enrichment analysis of GJ specific networks were performed using the g:GOSt tool in g:Profiler, using the Gene Ontology (GO: molecular function, and cellular components) (Reimand et al. 2007)

### SDS-PAGE and Western Blot Analysis of ANXA2 and Cx43

Whole-cell lysates (WCL) were prepared in RIPA buffer (150 mM NaCl, 1% Triton X-100, 0.5% sodium deoxycholate, 0.1% SDS, 50 mM Tris, pH 8.0, 1 mM PMSF). Proteins were separated on 10% SDS-PAGE gels and transferred to PVDF membranes (30 V, 3 h, 4°C). Membranes were blocked in 5% milk/TBST and incubated overnight at 4°C with primary antibodies: ANXA2 (Cell Signaling, 1:1000), Cx43 (Cell Signaling, 1:1000), GAPDH (Cell Signaling, 1:1000), or MMP3 (Proteintech, 1:1000). After TBST washes, membranes were incubated with HRP-conjugated secondary antibodies (1:2000) and developed using ECL. Membranes were stripped (0.2 M glycine, 0.1% SDS, 1% Tween-20, pH 2.2) and reprobed as needed.

### Immunofluorescence of Connexin 43 and Annexin A2

C6 and C6-13 cells were seeded in 8-well chamber slides and grown to ∼70% confluency. Cells were fixed with 4% paraformaldehyde (15 min), permeabilized with 0.2% Triton X-100 (5 min), and blocked with 2% BSA (30 min). Slides were incubated overnight at 4°C with mouse anti-ANXA2 (Proteintech) and rabbit anti-Cx43 (Sigma-Aldrich) in 1% BSA/0.1% Triton X-100. After PBS washes, cells were incubated with Alexa Fluor-conjugated secondary antibodies (goat anti-rabbit (550 nm) or goat anti-mouse (488 nm)) for 1 h at room temperature. Nuclei were counterstained with Hoechst. Images were acquired at 64× magnification and analyzed in ImageJ.

### Annexin A2-S100A10 Peptide Inhibition Zymography

In 6-well culture plates, confluent C6-13 cells were treated with 100 µM N-terminal ANXA2 peptides (Peptide-1: STVHEILCKLSL; Peptide-2: STVHEILCKLSLEG) or controls (scrambled peptide, DMSO). Peptides were obtained commercially (Peptide2.0, USA). After 24 h in serum-free medium, secretomes were concentrated using 30 kDa MWCO filters and normalized by BCA assay. Samples (50 µg protein) were resolved on 10% SDS-PAGE gels containing 0.1% gelatin under non-reducing conditions. Gels were washed in Triton X-100 renaturation buffer, incubated in developing buffer (50 mM Tris-HCl, pH 7.5, 5 mM CaCl₂, 1 µM ZnCl₂) for 24 h at 37°C, stained with Coomassie G-250, and destained. Proteolytic bands were quantified using ImageJ.

### Inhibition of 3D Tumor Invasion by N-terminal Annexin A2 peptides

C6-13 cells were seeded at 2 × 10⁴ cells/mL in ultra-low attachment 96-well plates and allowed to form spheroids over 4 days. Medium was replaced with basement membrane extract (BME; PathClear) and incubated for 1 h at 37°C. Spheroids were treated with ANXA2 Peptide-1 (50 µM), scrambled peptide (50 µM), DMSO (vehicle), or left untreated. Images were acquired at 0, 24, 48, 72, and 96 h at 10× magnification. Invasion was quantified by measuring spheroid area in ImageJ.

### Statistical Analysis

All experiments were performed with ≥3 biological replicates. Statistical significance was assessed using two-tailed Student’s tests. Differences between means were identified as statistically significant when *p < 0.05, **p < 0.005

## Results

### GJ Interactome In High Motility Glioma Reveals ANXA2 and ECM Regulators

Building on our previous identification of MMP upregulation in the secretome of C6-13 (Cx43-overexpressing) glioma cells (Aftab et al. 2019), we sought to identify proteins specifically recruited to GJ nexus that could link Cx43 to ECM activity. To do this we leveraged our previously published global Cx43 protein-protein interactome (Chen et al. 2012) and cross-referenced it against a GJ-enriched plasma membrane proteome generated from C6-13 glioma cells that identified 1737 at a 1% FDR across three or more of the 5 biological replicates. We observed a number of GJ-binding proteins, including actin, calmodulin (Duffy et al. 2007), caveolin-1 (Schubert et al. 2002), cadherin (Singh et al. 2005), drebrin (Butkevich et al. 2016), EPS15 (Girón-Pérez et al. 2011), HRS/HGS (Catarino et al. 2011), NEDD4 (Leykauf et al. 2006), TSG101, tubulin (Giepmans et al. 2001), UBC (Leithe et al. 2004), 14-3-3 zeta (Leithe et al. 2004), and ZO-1 (Giepmans et al. 2001), validating the specificity of our enrichment. Of the 110 proteins in the Cx43 interactome, 74 (67%) were similarly enriched in the GJ plasma membrane. Gene ontology analysis revealed significant enrichment of proteins involved in junctional assembly and ECM remodeling, suggesting that the GJ nexus functions as a scaffold for invasion-associated signaling networks. Amongst these proteins, we identified Annexin A2 (ANXA2), a known promoter of extracellular protease activity and that is frequently upregulated in GBM (Gao et al. 2013; Lee et al. 2004; Reeves et al. 1992) (Figure 1A-C). We further noted the presence of ANXA2 signaling protein S100A10 (Bharadwaj et al. 2021) suggesting the presence of an intact AnxA2/S100A10 complex at the GJ (Figure 1C).

**Figure 1:**
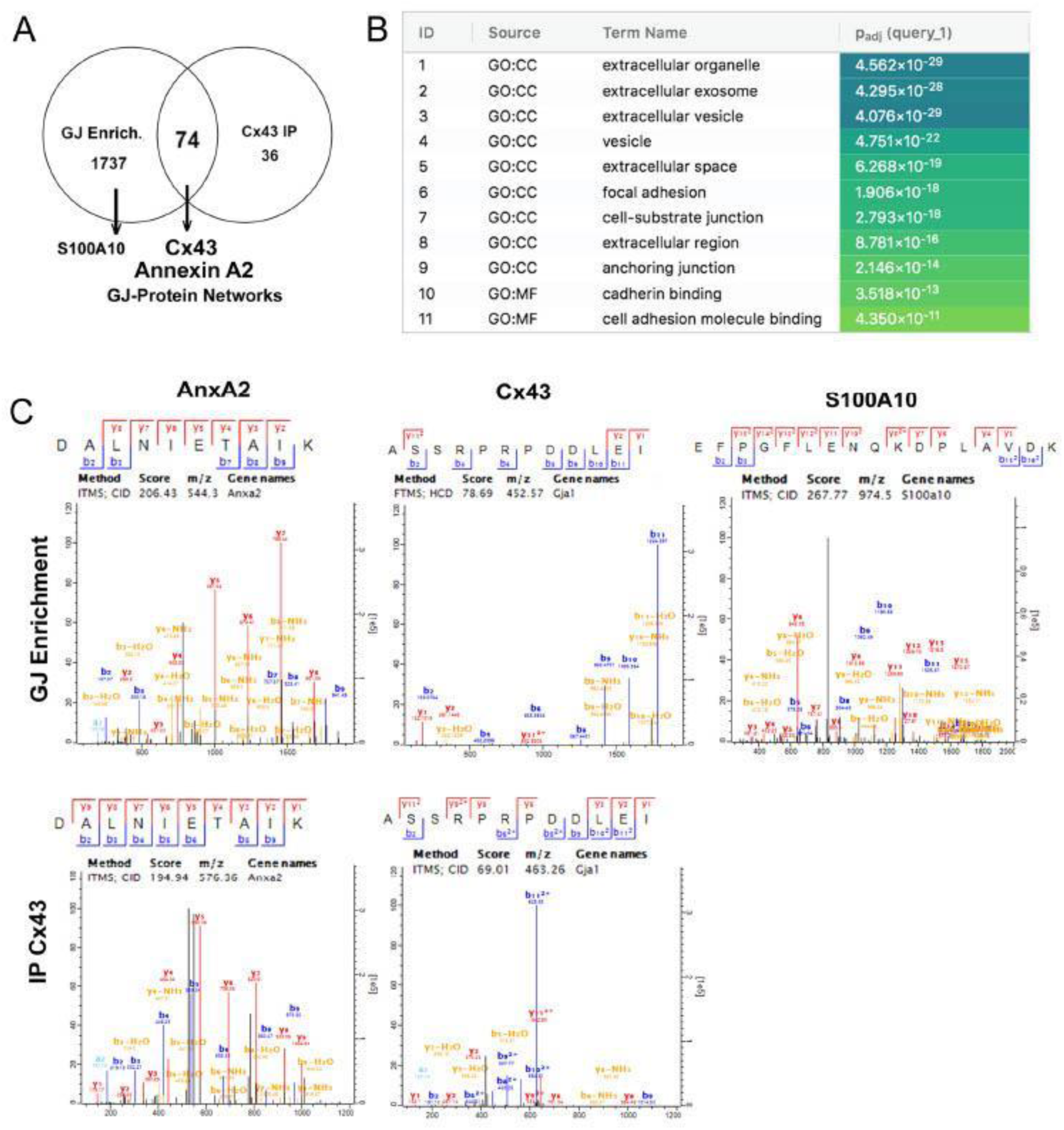
Integrated proteomic analysis of the Cx43 interactome identifies ANXA2 and ECM regulatory proteins at the gap junction. (A) Cross-referencing the published Cx43 interactome (Chen et al., 2012) with a gap junction (GJ)-enriched plasma membrane proteome shows that 67% of Cx43-associated proteins are similarly enriched at GJ membranes. (B) Gene ontology analysis reveals significant enrichment of proteins involved in junctional assembly and extracellular matrix (ECM) remodeling. (C) ANXA2 and Cx43 peptides were detected in GJ-enriched membrane fractions and Cx43 co-immunoprecipitation (Co-IP). The ANXA2 binding partner S100A10 was co-detected in GJ-enriched fractions, indicating the presence of an intact ANXA2/S100A10 complex at gap junctions.

### Cx43 interacts with ANXA2 and drives Its Recruitment to Gap Junctions

To validate the Cx43/ANXA2 interaction, reciprocal co-immunoprecipitation of C6-13 WCL using anti-ANXA2 antibodies as bait confirmed the Cx43/ANXA2 interaction, as evidenced by detection of Cx43 bands at ∼37–45 kDa via SDS-PAGE/Western Blot (Figure 2A). Comparison of whole cell lysates and GJ-containing membrane preparations confirmed successful GJ isolation, as demonstrated by strong Cx43 enrichment of Cx43 in the membrane fraction relative to whole cell lysates (Figure 2B) (Chen et al. 2012). To determine whether Cx43/ANXA2 interaction drives recruitment of ANXA2 to the plasma membrane, we compared ANXA2 levels in whole cell lysates and plasma membrane-enriched fractions from wild-type C6 and C6-13 cells. Although total ANXA2 levels were not statistically different between cell lines (Figures 2C-2D), ANXA2 was significantly enriched in plasma membrane fractions of C6-13 cells (Figures 2E-F). These data indicate that elevated Cx43 expression drives redistribution of ANXA2 to the plasma membrane rather than increasing total protein abundance.

**Figure 2:**
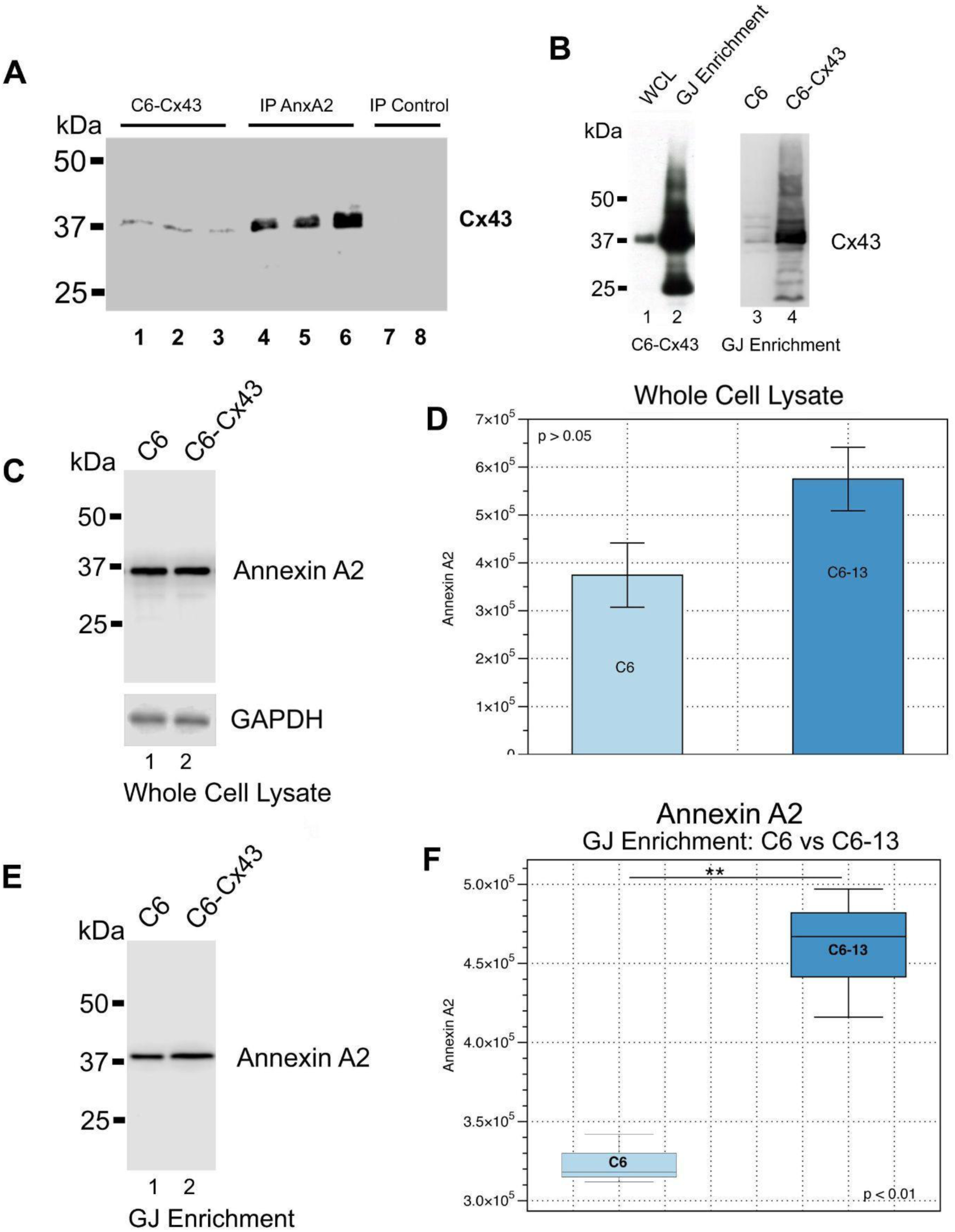
Biochemical validation of the Cx43/ANXA2 interaction and ANXA2 membrane redistribution in Cx43-overexpressing glioma cells. (A) Reciprocal Co-IP of C6-13 whole-cell lysates using anti-ANXA2 antibodies confirms the Cx43–ANXA2 interaction, with Cx43 detected at ∼37–45 kDa by SDS-PAGE/Western blot. (B) Western blot analysis shows strong enrichment of Cx43 in GJ-enriched membrane fractions relative to whole-cell lysates. (C–D) Total ANXA2 levels do not differ significantly between C6 and C6-13 whole-cell lysates (p > 0.05). (E–F) ANXA2 is significantly enriched in GJ-enriched membrane fractions from C6-13 cells compared with C6 controls, demonstrating Cx43-driven redistribution of ANXA2 to the plasma membrane.

### Cx43-overexpression Promotes ANXA2 Consolidation at Gap Junction Plaques

To characterize the subcellular distributions of Cx43 and ANXA2, we performed double immunofluorescence labeling in C6 and C6-13 glioma cell cultures and examined colocalization by confocal microscopy. In wild-type C6 cells, ANXA2 displayed diffuse cytoplasmic and perinuclear distribution with limited overlap with Cx43 (Figure 3A1-3). In contrast, C6-13 cells exhibited a pronounced redistribution of ANXA2 to sites of Cx43 enrichment across the plasma membrane. The merged images revealed increased yellow signal, indicating enhanced colocalization. (Figures 3B1-3). ANXA2 colocalization tracked local Cx43 expression level in both cell lines, occurring at the sparse Cx43 puncta present in C6 cells and more extensively in C6-13 cells, which display more numerous and more intense Cx43 puncta. This strong colocalization is consistent with the ANXA2 membrane enrichment observed biochemically. High-magnification insert further highlighted the consolidation of ANXA2 at Cx43-positive GJ plaques (Figures 3C-D). Together, these results support a model in which Cx43 drives local accumulation of ANXA2 at sites of gap junction formation.

**Figure 3:**
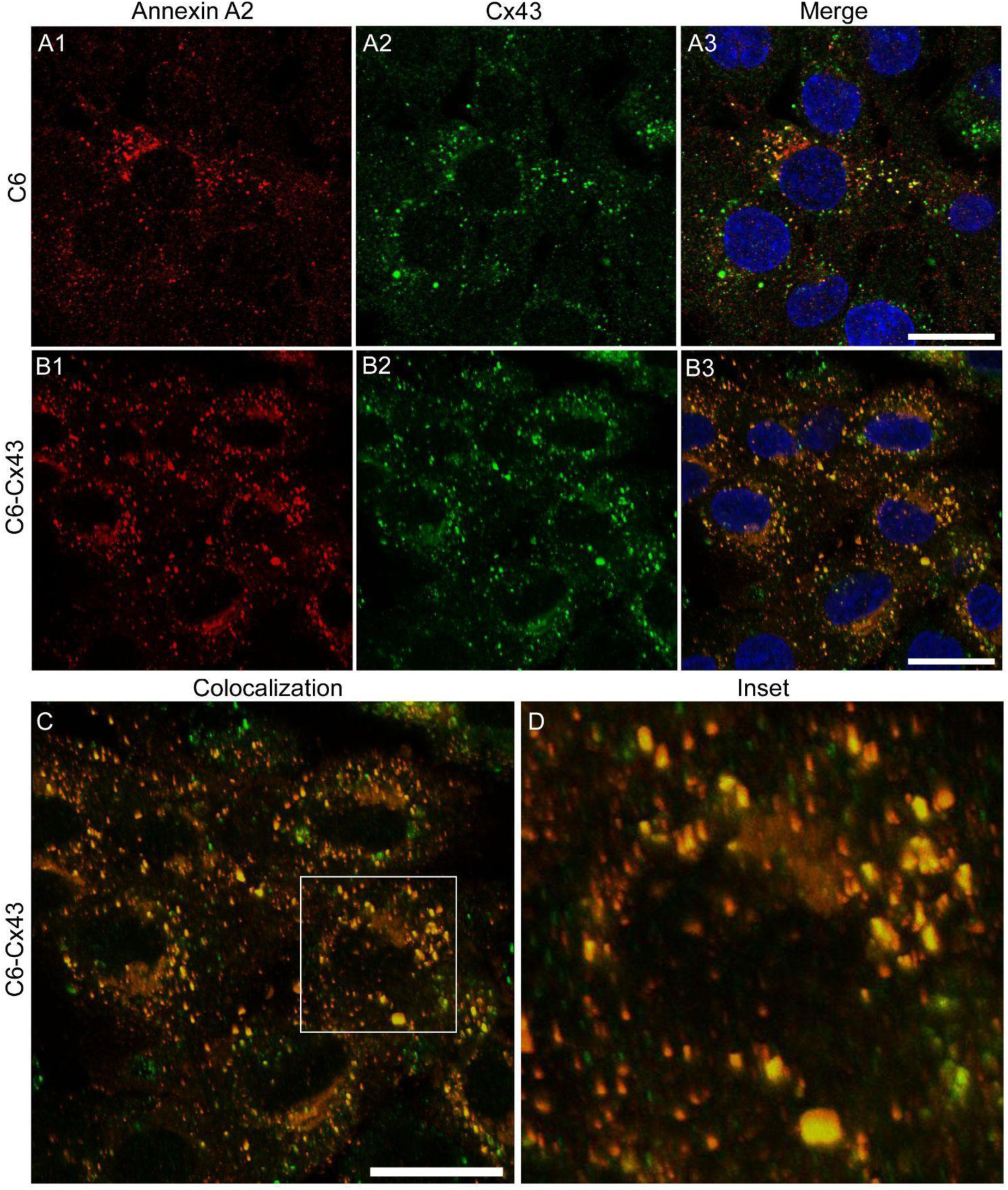
Confocal immunofluorescence confirms co-localization of Cx43 and ANXA2 at gap junction plaques in Cx43-overexpressing glioma cells. (A1–A3) In wild-type C6 cells, ANXA2 (red) displays diffuse cytoplasmic and perinuclear localization with limited overlap with Cx43 (green). Merged images show minimal yellow signal, indicating sparse colocalization. Nuclei are stained with Hoechst (blue). (B1–B3) In C6-13 cells, ANXA2 redistributes to the plasma membrane and strongly colocalizes with Cx43, producing increased yellow signal in merged images. (C) Merged Cx43/ANXA2 image (nuclear channel omitted) highlights membrane-localized colocalization. (D) High-magnification inset from panel C shows discrete yellow puncta at cell–cell interfaces, confirming consolidation of ANXA2 at Cx43-positive gap junction plaques.

### ANXA2 Mimetic Peptides Targeting the S100A10 Reduce MMP3 Activity

Having established that ANXA2 and S100A10 are co-enriched in GJ membrane fractions and that elevated Cx43 expression drives ANXA2 redistribution to GJ plaques, we next sought to determine whether disruption of the ANXA2/S100A10 complex could alter MMP3 activity. The binding of S100A10 to the N-terminal domain of ANXA2 is known to facilitate plasmin production and downstream MMP activation (Bolon et al. 2004; Sharma and Sharma 2007), making this interaction an attractive target for pharmacological intervention. The crystal structure of the ANXA2/S100A10 complex (RCSB PDB: 5LPU)(Figure 4A) reveals the key residues of the N-terminal S100A10-binding domain, including GLY15, GLU14, SER12, LEU11, CYS9, and HIS5 (Figure 4B). Based on this structural data, two ANXA2-mimetic peptides; Peptide 1 (STVHEILCKLSL) and Peptide 2 (STVHEILCKLSLEG) were designed to competitively inhibit S100A10 binding at this interface, alongside a scrambled sequence control (HSLVLKEGILTSCE, Figure 4C).

**Figure 4:**
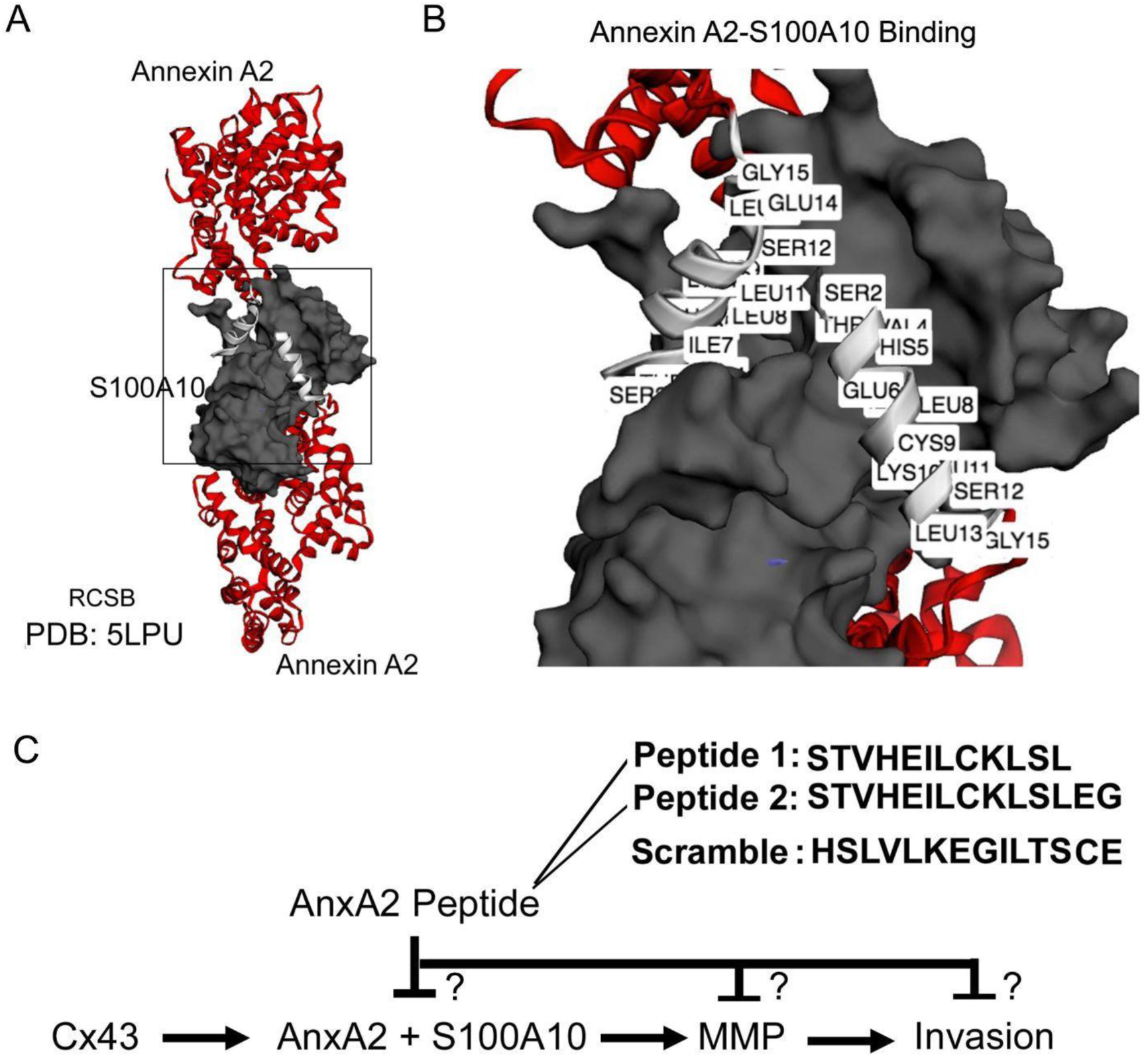
Structural basis for ANXA2-mimetic peptide design targeting the ANXA2/S100A10 binding interface. (A) Three-dimensional structure of the ANXA2/S100A10 heterotetramer (PDB: 5LPU), with S100A10 shown in grey. (B) High-resolution view of the ANXA2 N-terminal S100A10-binding domain, highlighting key interface residues (HIS5, CYS9, LEU11, SER12, GLU14, GLY15). (C) Proposed Cx43-mediated invasion pathway involving ANXA2/S100A10-dependent activation of MMP3. Peptide-1 (STVHEILCKLSL) and Peptide-2 (STVHEILCKLSLEG) were designed to competitively inhibit the S100A10-binding interface of ANXA2. The scrambled control peptide (HSLVLKEGILTSCE) contains the same amino acids as Peptide-2 in randomized order.

To assess the effects of ANXA2/S100A10 disruption on MMP3 activity, secretomes from C6-13 cells treated with Peptide 1, Peptide 2, scrambled control, or vehicle (DMSO) were analyzed by gelatin zymography (Figure 5A). MMP3 was detected as a band at ∼60 kDa, consistent with its known molecular weight. Quantification of band signal intensity revealed that cells treated with Peptide 1 and Peptide 2 showed an ∼50% reduction in MMP3 activity relative to untreated cells (p < 0.05) (Figure 5B). In contrast, scrambled peptide-treated cells showed no significant reduction in MMP3 activity relative to untreated controls (Figure 5B). Together, these results demonstrate that the ANXA2/S100A10 complex is a critical upstream regulator of MMP3 activation in Cx43-overexpressing glioma cells.

**Figure 5:**
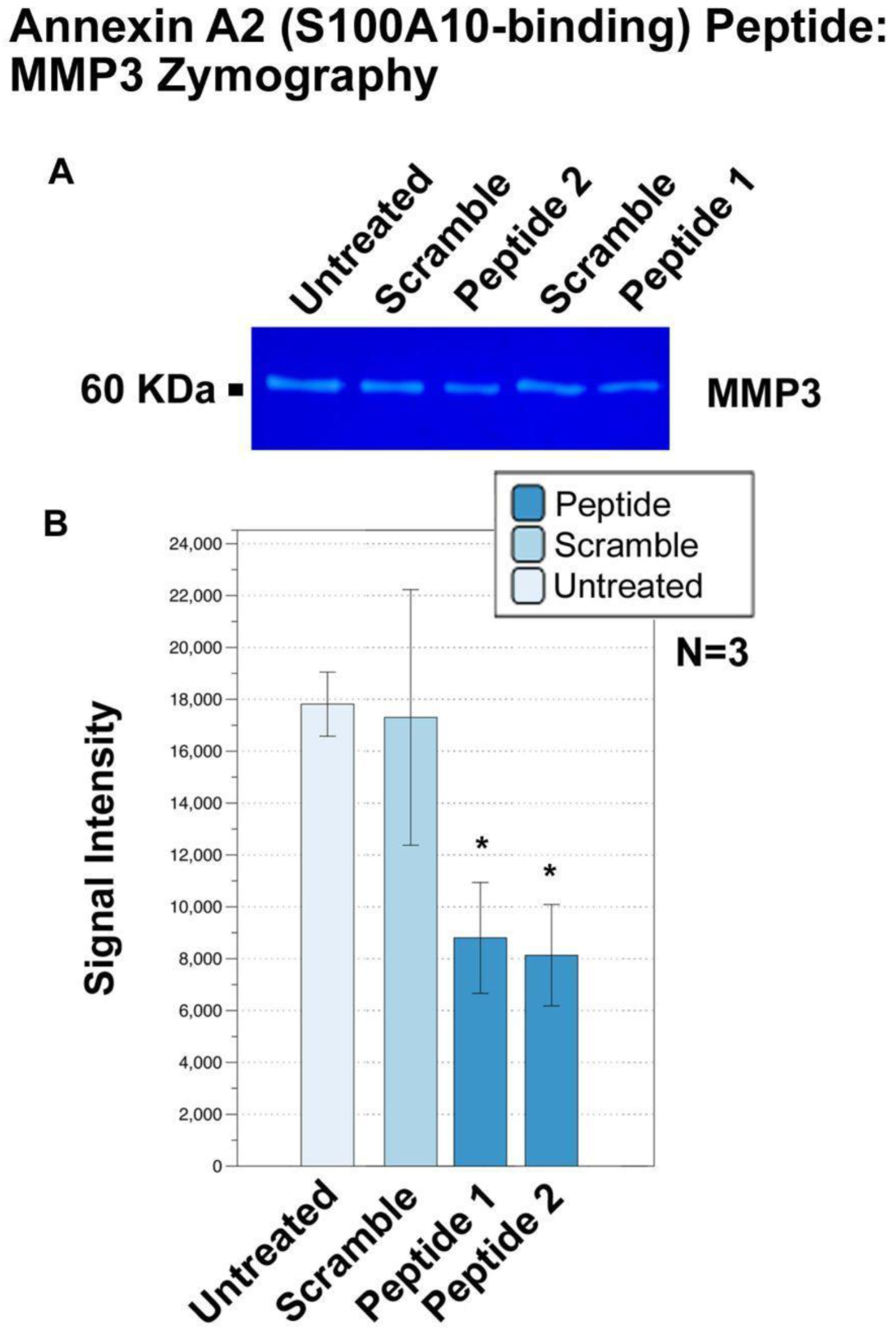
ANXA2-mimetic peptides targeting the S100A10-binding domain reduce MMP3 activity in Cx43-overexpressing glioma cells. (A) Gelatin zymography of secretomes from C6-13 cells treated with ANXA2 Peptide-1, Peptide-2, scrambled peptide, or left untreated. MMP3 appears as a ∼60 kDa band. (B) Quantification of zymography band intensities (N = 3). Peptide-1 and Peptide-2 reduce MMP3 activity by ∼50% relative to untreated controls (p < 0.05). Scrambled peptide shows no significant effect, confirming specificity of ANXA2/S100A10 disruption.

### ANXA2/S100A10 Disruption Reduces Glioma Invasion in a 3D Spheroids

To determine whether reduced MMP3 activity translates to functional inhibition of invasion, C6-13 spheroids were embedded in basement membrane extract (BME) (PathClear) and treated with ANXA2 Peptide-1, scrambled peptide, DMSO, or left untreated. Invasion was monitored by imaging at 0, 24, 48, 72, and 96 hours and quantified by measuring the increase in spheroid area relative to the 24-hour baseline (Figure 6A-B). Quantification of spheroid area revealed a significant reduction in invasion beginning at 48 hours and persisting through 96 hours, with Peptide-1 producing an approximately 30% decrease in invasive area relative to controls (Figure 6C–D). No significant differences were observed between untreated, vehicle, and scrambled peptide conditions, confirming specificity of the inhibitory effect. Taken together, these results demonstrate that targeted disruption of the ANXA2/S100A10 complex using N-terminal mimetic peptides significantly suppresses glioma invasion into the ECM, an effect attributable to the downstream reduction in MMP3 activity observed in preceding experiments.

**Figure 6:**
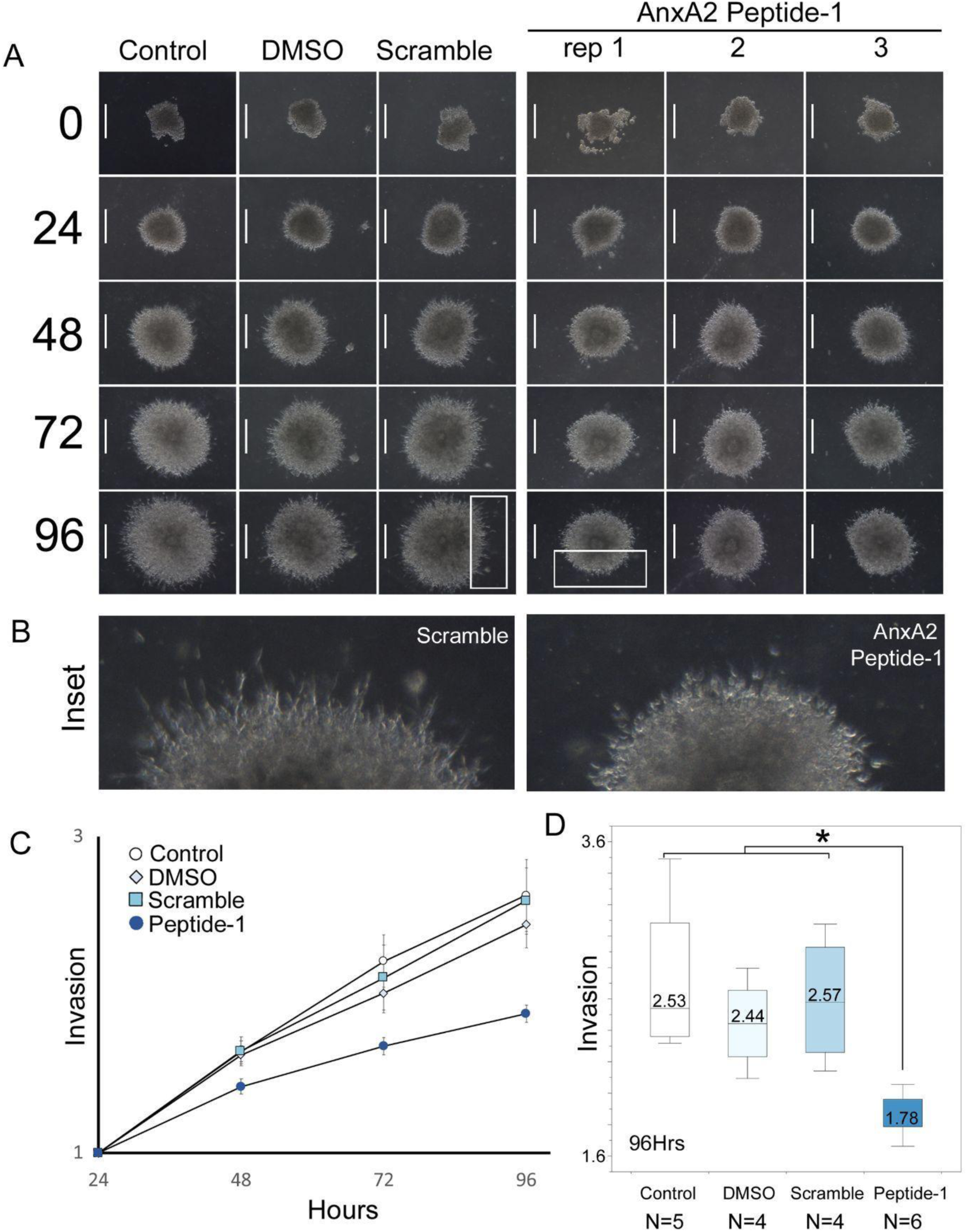
Competitive inhibition of the ANXA2/S100A10 complex reduces glioma spheroid invasion into a 3D ECM-replicating matrix. (A) Representative brightfield images of C6-13 spheroids embedded in basement membrane extract (BME) and treated with ANXA2 Peptide-1, scrambled peptide, DMSO, or left untreated, imaged over 0–96 hours. (B) High-magnification images at 96 hours show prominent invasive protrusions in scrambled peptide-treated spheroids and markedly reduced protrusions in Peptide-1-treated spheroids. (C) Quantification of spheroid invasion over time, expressed as fold-increase in spheroid area relative to the 24-hour baseline. Peptide-1 significantly reduces invasion beginning at 48 hours. (D) Boxplot of invasion at 96 hours (Control N=5, DMSO N=4, Scramble N=4, Peptide-1 N=6). Peptide-1-treated spheroids show significantly reduced invasive area (1.78-fold) compared with untreated (2.53-fold), DMSO (2.44-fold), and scrambled peptide (2.57-fold) controls (p < 0.05). No significant differences are observed among control conditions.

## Discussion

Glioblastoma (GBM) remains one of the most treatment-refractory malignancies, with its lethality driven by the tumor’s drug resistance and the capacity of tumor cells to invade the surrounding CNS and evade surgical resection (Giese and Westphal 1996). The gap junction protein Cx43 has emerged as a paradoxical driver of this invasive phenotype. While early studies established Cx43 as a tumor suppressor on the basis of its ability to slow glioma cell proliferation (Fu et al. 2004; Huang et al. 1998; Zhu et al. 1991), accumulating evidence has since demonstrated that Cx43 actively promotes glioma invasion, migration, and resistance to temozolomide (TMZ) chemotherapy (Aftab et al. 2019; Naus et al. 2016; Sin et al. 2012). Cx43-driven invasion has been observed across multiple *in vitro* and *ex vivo* models and has been linked to dysregulation of ECM-associated proteins, including MMP2, MMP3, osteopontin, CCN1, and CCN3/NOV (Bates et al. 2007; Naus et al. 2000; Oliveira et al. 2005; Sin et al. 2008; Zhang et al. 2003). However, the upstream molecular mechanisms linking Cx43 to ECM-remodeling remain poorly understood, making their characterization a key priority for developing novel therapeutic strategies.

In this study, we identify and functionally validate a previously uncharacterized Cx43→ANXA2→S100A10→MMP3 signaling axis that directly links gap junction signaling to ECM proteolysis. Building on our previous identification of MMP3 upregulation in the secretome of Cx43-overexpressing glioma cells (Aftab et al. 2019), this study employed a multi-proteomic approach to identify the molecular intermediaries linking Cx43 to ECM protease activity. Cross-referencing our previously published global Cx43 interactome (Chen et al. 2012) against a new GJ-enriched plasma membrane proteome revealed that 67% of the Cx43 interactome (74 of 110 proteins) was similarly enriched at GJ membranes (Figure 1A). Gene ontology analysis of this network demonstrated significant enrichment of factors underlying both junctional assembly and ECM remodeling, providing the first high-throughput proteomic evidence that the GJ signaling nexus actively recruits ECM regulatory machinery (Figure 1B).

Amongst these, Annexin A2 (ANXA2) emerged as a compelling candidate due to its established role in plasmin generation, MMP activation, and its frequent upregulation in aggressive GBM (Figure 1C) (Reeves et al. 1992; Zhai et al. 2011).

Biochemical validation confirmed the Cx43/ANXA2 interaction and revealed an important mechanistic detail that while total ANXA2 levels were statistically indistinguishable between wild-type C6 and C6-13 cells, ANXA2 was found to be significantly enriched in GJ containing plasma membrane fractions of Cx43-overexpressing cells (Figure 2). These findings indicate that Cx43 drives membrane redistribution of ANXA2 rather than transcriptional upregulation of total protein, suggesting that the GJ nexus may functions as an organizing platform to recruit regulators of invasion. Confocal immunofluorescence confirmed this redistribution, showing consolidation of ANXA2 at Cx43-positive GJ plaques in C6-13 cells that was largely absent in wild-type C6 cells (Figure 3). The co-enrichment of S100A10 alongside ANXA2 in GJ–enriched membrane fractions lend further credibility of intact ANXA2/S100A10 complexes associated to Cx43 at GJ (Figure 1C). ANXA2/S100A10 is recognized as a key oncogenic plasminogen receptor, functions to concentrate plasminogen and tissue plasminogen activator (tPA), thereby generating plasmin and activating downstream MMPs, including MMP3 (Madureira et al. 2021, Waisman et al. 2007) (Figure 4). To test whether this complex mediates Cx43-driven MMP3 activity, we designed N-terminal ANXA2 mimetic peptides that competitively inhibit S100A10 binding. Disruption of the ANXA2/S100A10 interface resulted in a ∼50% reduction in MMP3 activity and expression, demonstrating that this complex is a critical upstream regulator of MMP3 in Cx43-overexpressing glioma cells (Figure 5). The functional significance of this pathway was confirmed using a 3D spheroid invasion assay. The 3D spheroid model provides a methodological advantage over conventional 2D invasion assays, as the BME matrix more faithfully recapitulates the biochemical and mechanical properties of the in vivo ECM (Schiffer et al. 2018; Vinci, Box, and Eccles 2015). Here, C6-13 spheroids suspended in basement membrane extract (BME) treated with ANXA2 Peptide-1 showed a significant reduction in invasive area relative to untreated, DMSO, and scrambled peptide controls. These findings confirm that inhibition of the ANXA2/S100A10 complex attenuates Cx43-driven ECM remodeling (Figure 6). Collectively, these findings identity the ANXA2/S100A10 complex as a functional downstream effector of Cx43-driven invasion, consistent with prior evidence linking ANXA2 to glioma aggressiveness and elevated MMP activity to poor GBM prognosis (Forsyth et al. 1999; Xue et al. 2017; Zhai et al. 2011).

Together, the data presented here define a novel Cx43→ANXA2→S100A10→MMP3 signaling axis that links gap junction biology to ECM remodeling and glioma invasion, identifying ANXA2 as the key molecular intermediary connecting Cx43 to protease activation within the extracellular space (Sin, Crespin, and Mesnil 2012; Naus, Aftab, and Sin 2016). By establishing the ANXA2/S100A10 complex as a functional downstream effector of Cx43-driven invasion, this work significantly advances our mechanistic understanding of how Cx43 contributes to glioblastoma malignancy. Moreover, the ability of N-terminal ANXA2 mimetic peptides to markedly reduce both MMP3 activity and 3D spheroid invasion highlights the ANXA2/S100A10 complex as a promising therapeutic target for disrupting Cx43-mediated ECM remodeling and limiting glioma progression.

## ACKNOWLEDGEMENTS

CG was supported by an NSERC-USRA, Dr. Bruce and Mrs. Jane Forrest Memorial Scholarship. VCC was supported by the National Science and Engineering Research Council of Canada (NSERC, RGPIN-2018-04261), Canadian Foundation for Innovation (CFI), BC Knowledge Development Fund, and the Brandon University Research Committee (BURC).

## AUTHOR CONTRIBUTIONS

Conceptualization, VCC, LJF, CCN; Formal Analysis CG, EO, VCC; Funding Acquisition, VCC, CCN, Methodology, EO, CG, VCC, WCS; Project Administration, VCC; Supervision; VCC, CCN, LJF.; Writing CG, VCC.

## References

Aftab, Qurratulain, Marc Mesnil, Emmanuel Ojefua, Alisha Poole, Jenna Noordenbos, Pierre-Olivier Strale, Chris Sitko, et al. 2019. “Cx43-Associated Secretome and Interactome Reveal Synergistic Mechanisms for Glioma Migration and MMP3 Activation.” Frontiers in Neuroscience 13: 143.

Bates, Dave C., W. C. Sin, Q. Aftab, and C. C. Naus. 2007. “Connexin43 Enhances Glioma Invasion by a Mechanism Involving the Carboxy Terminus.” Glia 55 (15): 1554–64.

Behrens, J., J. F. Bechberger, and C. C. Naus. 2010. “The Connexin43 Carboxy-Terminus Controls Surface Protease Scaffolding during Glioma Cell Dispersion.” Journal of Neuro-Oncology 98 (2): 171–181.

Bellail, A. C., S. B. Hunter, D. T. Brat, C. Tan, and O. Olson. 2004. “Microenvironmental Mechanisms of Glioblastoma Invasion.” The International Journal of Biochemistry & Cell Biology 36 (6): 1055–1069.

Bharadwaj, Anupama, Andrew M. G. Cochrane, and David M. Waisman. 2021. “S100A10: A Multifunctional Scaffold Protein in Human Health and Disease.” Biomolecules 11 (12): 1849.

Bolon, Isabelle, Hong-Ming Zhou, Yves Charron, Annelise Wohlwend, and Jean-Dominique Vassalli. 2004. “Plasminogen Mediates the Pathological Effects of Urokinase-Type Plasminogen Activator Overexpression.” The American Journal of Pathology 164 (6): 2299–2304.

Butkevich, Eugenia, Benjamin Bodin, Michael S. S. Sharrow, et al. 2016. “Drebrin Regulates Connexin43 Trafficking and Gap Junction Assembly in Astrocytes.” PLOS ONE 11 (6): e0157073.

Cabral-Pacheco, S. M., I. Araujo, and C. Cabral. 2020. “The Role of Matrix Metalloproteinases in Glioma Progression and Extracellular Matrix Remodeling.” Cancers 12 (5): 1120.

Catarino, Sandra, Inês Marques, and Henrique Girão. 2011. “The Ubiquitin-Protein Ligase NEDD4-1 Regulates Connexin43 Localization and Degradation.” Journal of Cell Science 124 (Pt 15): 2580– 2591.

Cawston, T. E., and D. A. Young. 2010. “Matrix Metalloproteinases and Their Inhibitors in Connective Tissue Remodeling and Tumor Metastasis.” British Journal of Pharmacology 161 (7): 1421–1434.

Chen, Ricky, Matthew Smith-Cohn, Adam L. Cohen, and Howard Colman. 2017. “Glioma Subclassifications and Their Clinical Significance.” Neurotherapeutics: The Journal of the American Society for Experimental NeuroTherapeutics 14 (2): 284–97.

Chen, Vincent C., Anders R. Kristensen, Leonard J. Foster, and Christian C. Naus. 2012. “Association of Connexin43 with E3 Ubiquitin Ligase TRIM21 Reveals a Mechanism for Gap Junction Phosphodegron Control.” Journal of Proteome Research 11 (12): 6134–46.

Cina, C., C. Onzon, and Christian C. Naus. 2009. “The Cytoplasmic Domain of Connexin43 Modulates Astrocyte Adhesivity and Inward Rectification.” Journal of Cell Science 122 (15): 2691–2701.

Cooper, Lee A. D., David A. Gutman, Chih-Wei Chang, Sharath R. Chisolm, Carlos S. Moreno, David T. Brat, and Jun Kong. 2012. “An Integrative Approach for Profiling the In Situ Cellular Composition of Glioblastoma.” The American Journal of Pathology 180 (3): 908–919.

Demuth, Tim, and Michael E. Berens. 2004. “Molecular Mechanisms of Glioma Cell Migration and Invasion.” Journal of Neuro-Oncology 70 (2): 217–228.

Dong, Hui, Xing-Wang Zhou, Xiang Wang, Yuan Yang, Jie-Wen Luo, Yan-Hui Liu, and Qing Mao. 2017. “Complex Role of Connexin 43 in Astrocytic Tumors and Possible Promotion of Glioma-associated Epileptic Discharge (Review).” Molecular Medicine Reports 16 (6): 7890–7900.

Duffy, Heather S., John J. S. Sorgen, and David C. Spray. 2007. “Calmodulin Binding to the Connexin43 Carboxy Terminus Modulates Gap Junction Gating.” Journal of Biological Chemistry 282 (51): 37190–37199.

Duffy, Heather S., S. M. Ashton, and David C. Spray. 2002. “The Supramolecular Gap Junction Nexus: Protein Complexes Associated with Connexin Channels.” Glia 40 (2): 121–132.

Forsyth, P. A., H. Wong, T. D. Laing, N. B. Rewcastle, D. G. Morris, H. Muzik, K. J. Leco, et al. 1999. “Gelatinase-A (MMP-2), Gelatinase-B (MMP-9) and Membrane Type Matrix Metalloproteinase-1 (MT1-MMP) Are Involved in Different Aspects of the Pathophysiology of Malignant Gliomas.” British Journal of Cancer 79 (11-12): 1828–35.

Fu, Christine T., John F. Bechberger, Mark A. Ozog, Bernard Perbal, and Christian C. Naus. 2004. “CCN3 (NOV) Interacts with connexin43 in C6 Glioma Cells: Possible Mechanism of Connexin-Mediated Growth Suppression.” The Journal of Biological Chemistry 279 (35): 36943–50.

Furnari, Frank B., Tim Fenton, Robert M. Bachoo, Akitake Mukasa, Jayne M. Stommel, Alexander Stegh, William C. Hahn, et al. 2007. “Malignant Astrocytic Glioma: Genetics, Biology, and Paths to Treatment.” Genes & Development 21 (21): 2683–2710.

Gao, L., X. Zhang, and J. Liu. 2013. “Annexin A2 Dysregulation Drives Aberrant Extracellular Proteolysis in High-Grade Glioblastoma.” Oncology Reports 29 (4): 1501–1509.

Giepmans, Kees B. G., Wouter H. Moolenaar, and Wun-Chey Sin. 2001. “Connexin43 Interacts directly with ZO-1 and Tubulin to Scaffold the Subplasmalemmal Cytoskeleton.” Current Biology 11 (17): 1364–1368.

Gielen, Paul R., Qurratulain Aftab, Noreen Ma, Vincent C. Chen, Xiaoting Hong, Shannon Lozinsky, Christian C. Naus, and Wun Chey Sin. 2013. “Connexin43 Confers Temozolomide Resistance in Human Glioma Cells by Modulating the Mitochondrial Apoptosis Pathway.” Neuropharmacology 75 (December): 539–48.

Giese, A., and M. Westphal. 1996. “Glioma Invasion in the Central Nervous System.” Neurosurgery 39 (2): 235–50.

Girón-Pérez, Daniel A., Elizabeth S. S. EPS15 Working Group, et al. 2011. “EPS15 Coordinates Internalization Dynamics of Intercellular Plaques.” Biochemical Journal 438 (2): 321–331.

Goodenough, D. A., and D. L. Paul. 2009. “Gap Junctions.” Cold Spring Harbor Perspectives in Biology 1 (1): a002576.

Hervé, J. C., N. Bourmeyster, D. Sarrouilhe, and H. S. Duffy. 2007. “Protein–Protein Interactions with Connexin43: Regulation and Function.” Progress in Biophysics and Molecular Biology 94 (1-2): 120–143.

Holland, E. C. 2000. “Glioblastoma Multiforme: The Clinical Problem and Molecular Changes.” Oncogene 19 (53): 6100–6107.

Huang, R. P., Y. Fan, M. Z. Hossain, A. Peng, Z. L. Zeng, and A. L. Boynton. 1998. “Reversion of the Neoplastic Phenotype of Human Glioblastoma Cells by Connexin 43 (cx43).” Cancer Research 58 (22): 5089–96.

Iwadate, Yasuo. 2016. “Epithelial-Mesenchymal Transition in Glioblastoma Progression.” Oncology Letters 11 (3): 1615–20.

Kameritsch, Petra, Kristin Pogoda, and Ulrich Pohl. 2012. “Channel-Independent Influence of Connexin 43 on Cell Migration.” Biochimica et Biophysica Acta 1818 (8): 1993–2001.

Kapoor, C., S. Vaidya, and V. Kumar. 2016. “Upstream Regulators of Aberrant MMP Activity in the Glioblastoma Microenvironment.” Tumor Biology 37 (9): 11621–11635.

Laird, Dale W. 2010. “The Gap Junction Proteome and its Relationship to Disease.” Trends in Cell Biology 20 (2): 92–101.

Lee, I. H., A. M. Magliocco, and D. M. Waisman. 2004. “Annexin A2 Heterotetramer Enhances Plasminogen Activation on the Surface of Aggressive Brain Tumor Cells.” Journal of Biological Chemistry 279 (4): 2470–2478.

Leithe, Edward, Ingeborg S. S. YWHAZ Hub, and Kirsten Sandvig. 2004. “Ubiquitin-Dependent Endocytosis of Connexin43 Mediated by Interactions with 14-3-3 Zeta (YWHAZ) and UBC Complex.” Journal of Biological Chemistry 279 (31): 32450–32460.

Leykauf, Nicole, Martin S. S. Leykauf, and Christian C. Naus. 2006. “The Ubiquitin Ligase NEDD4 Interacts with Connexin43 to Mitigate High-Motility Membrane Pools.” Journal of Cell Science 119 (10): 2100–2109.

Liang, X., H. Wang, and Q. Mao. 2020. “The Non-Channel Scaffolding Intermediaries of Connexin43 in Tumor Pathology.” Cellular Signalling 68: 109520.

Madureira, Patricia A., David M. Waisman, et al. 2021. “The ANXA2/S100A10 Heterotetramer as a Key Oncogenic Plasminogen Receptor in Aggressive Malignancies.” Biomolecules 11 (12): 1772.

Machtaler, S., J. Bechberger, and Christian C. Naus. 2011. “The Carboxy-Terminal Domain of Connexin43 Scaffolds Cytoskeletal Anchors Essential for Glioblastoma Spreading.” Glia 59 (6): 932–944.

Mohiuddin, M., and H. Wakimoto. 2021. “Extracellular Matrix Remodeling in Glioblastoma Invasiveness and Therapeutic Resistance.” Frontiers in Oncology 11: 636544.

Munoz, J. L., V. Rodriguez-Cruz, S. J. Greco, S. H. Ramkissoon, K. L. Ligon, and P. Rameshwar. 2014. “Temozolomide Resistance in Glioblastoma Cells Occurs Partly through Epidermal Growth Factor Receptor-Mediated Induction of Connexin 43.” Cell Death & Disease 5 (March): e1145.

Nakada, M., S. Kita, and Y. Hayashi. 2007. “Roles of Matrix Metalloproteinases in Glioma Invasion.” Frontiers in Bioscience 12: 1205–1218.

Naus, Christian C., Qurratulain Aftab, and Wun Chey Sin. 2016. “Common Mechanisms Linking connexin43 to Neural Progenitor Cell Migration and Glioma Invasion.” Seminars in Cell & Developmental Biology 50 (February): 59–66.

Naus, Christian C., and Dale W. Laird. 2010. “Implication of Connexin Channels and Hemichannels in the Progression of Brain Malignancies.” Nature Reviews Cancer 10 (6): 435–441.

Oliveira, Roxane, Christo Christov, Jean Sébastien Guillamo, Sophie de Boüard, Stéphane Palfi, Laurent Venance, Marcienne Tardy, and Marc Peschanski. 2005. “Contribution of Gap Junctional Communication between Tumor Cells and Astroglia to the Invasion of the Brain Parenchyma by Human Glioblastomas.” BMC Cell Biology 6 (1): 7.

Ostrom, Quinn T., Haley Gittleman, Peter Liao, Chaturia Rouse, Yanwen Chen, Jacqueline Dowling, Yingli Wolinsky, Carol Kruchko, and Jill Barnholtz-Sloan. 2014. “CBTRUS Statistical Report: Primary Brain and Central Nervous System Tumors Diagnosed in the United States in 2007-2011.” Neuro-Oncology 16 Suppl 4 (October): iv1–63.

Parkin, D. M., F. Bray, J. Ferlay, and P. Pisani. 2001. “Estimating the World Cancer Burden: Globocan 2000.” International Journal of Cancer 94 (2): 153–56.

Plotkin, L. I., and T. Bellido. 2013. “Connexin43 Hemichannels and Gap Junction Plaques: Multi-Functional Platforms for Transmembrane and Intracellular Signaling.” Biochimica et Biophysica Acta (BBA) - Biomembranes 1828 (1): 120–128.

Pridham, Kevin J., et al. 2022. “Connexin43 Scaffolds Non-Channel Migration Machinery and Confers Front-Line Chemotherapeutic Resistance in Aggressive Glioblastoma Subtypes.” Oncogene 41 (14): 2011–2024.

Qin, L., X. Zhao, and G. G. Ghatnekar. 2016. “Structural Dynamics and Scaffolding Properties of Connexin Gap Junctional Arrays.” Journal of Biological Chemistry 291 (15): 7901–7912.

Reeves, S. A., C. Chavez-Kappel, R. Davis, M. Rosenblum, and M. A. Israel. 1992. “Developmental Regulation of Annexin II (Lipocortin 2) in Human Brain and Expression in High Grade Glioma.” Cancer Research 52 (24): 6871–76.

Reimand, Jüri, Meelis Kull, Hedi Peterson, Jaanus Hansen, and Jaak Vilo. 2007. “g:Profiler—a Web-Based Toolset for Functional Profiling of Gene Lists from Large-Scale Experiments.” Nucleic Acids Research 35 (Web Server issue): W193–W200.

Schiffer, Davide, Laura Annovazzi, Cristina Casalone, Cristiano Corona, and Marta Mellai. 2018. “Glioblastoma: Microenvironment and Niche Concept.” Cancers 11 (1): 35.

Schubert, Arthur L., Christian C. Naus, and Michael S. S. Lisanti. 2002. “Caveolin-1 (CAV1) Scaffolds Connexin43 Plasma Membrane Domains to Regulate Junctional Stability.” Biochemistry 41 (18): 5754–5764.

Sharma, Mahesh C., and Meena Sharma. 2007. “The Role of Annexin II in Angiogenesis and Tumor Progression: A Potential Therapeutic Target.” Current Pharmaceutical Design 13 (35): 3568–75.

Sharrow, Michael S. S., et al. 2008. “Connexin43 Carboxy-Terminal Structural Domains Coordinate Non-Channel Tumor Intracellular Signalling Cascades.” Cellular Signalling 20 (9): 1640–1651.

Sin, Wun Chey, John F. Bechberger, Walter J. Rushlow, and Christian C. Naus. 2008. “Dose-Dependent Differential Upregulation of CCN1/Cyr61 and CCN3/NOV by the Gap Junction Protein Connexin43 in Glioma Cells.” Journal of Cellular Biochemistry 103 (6): 1772–82.

Sin, Wun-Chey, Sophie Crespin, and Marc Mesnil. 2012. “Opposing Roles of connexin43 in Glioma Progression.” Biochimica et Biophysica Acta 1818 (8): 2058–67.

Singh, David B., Vincent C. Chen, and Chris S. S. Cadherin Group. 2005. “Co-Assembly of N-Cadherin and Connexin43 Junctional Scaffolds in Rat Glioma Models.” Journal of Biological Chemistry 280 (19): 18940–18949.

Sorgen, John J., et al. 2018. “Structural Variations within the Connexin43 C-Terminal Tail Dictate Intercellular Protein-Protein Assemblies.” Journal of Biological Chemistry 293 (22): 8401–8415.

Thévenin, N., et al. 2013. “The Cytoplasmic Carboxy-Terminal Tail (CT) of Connexin43 Coordinates Multiprotein Scaffolding Interactions Associated with Cell Migration.” Biochimica et Biophysica Acta 1828 (8): 1980–1992.

Waisman, David M., Anupama Bharadwaj, et al. 2007. “The Role of the S100A10/Annexin A2 Complex in Angiogenesis and Matrix Metalloproteinase Activation.” Current Pharmaceutical Design 13 (35): 3568–3575.

Weller, Michael, Wolfgang Wick, Ken Aldape, Michael Brada, Mitchell Berger, Stefan M. Pfister, Ryo Nishikawa, et al. 2015. “Glioma.” Nature Reviews. Disease Primers 1 (July): 15017.

Xue, Qiang, Li Cao, Xiao-Yan Chen, Jing Zhao, Liang Gao, San-Zhong Li, and Zhou Fei. 2017. “High Expression of MMP9 in Glioma Affects Cell Proliferation and Is Associated with Patient Survival Rates.” Oncology Letters 13 (3): 1325–30.

Yilmaz, M., and G. Christofori. 2009. “EMT, the Cytoskeleton, and Cancer Cell Invasion.” Cancer and Metastasis Reviews 28 (1-2): 15–33.

Zhai, Haiyan, Suchitra Acharya, Iordanis Gravanis, Saira Mehmood, Roberta J. Seidman, Kenneth R. Shroyer, Katherine A. Hajjar, and Stella E. Tsirka. 2011. “Annexin A2 Promotes Glioma Cell Invasion and Tumor Progression.” The Journal of Neuroscience 31 (40): 14346–60.

Zhang, Wei, Chiedozie Nwagwu, Duc Minh Le, V. Wee Yong, Hua Song, and William T. Couldwell. 2003. “Increased Invasive Capacity of connexin43-Overexpressing Malignant Glioma Cells.” Journal of Neurosurgery 99 (6): 1039–46.

Zhao, X., L. Qin, and R. G. Gourdie. 2015. “The Connexin43 Carboxy-Terminus Interaction Net: Scaffolding Intercellular Junction Complexes.” Cell Communication & Adhesion 22 (2): 45–56.

Zhu, D., S. Caveney, G. M. Kidder, and C. C. Naus. 1991. “Transfection of C6 Glioma Cells with Connexin 43 cDNA: Analysis of Expression, Intercellular Coupling, and Cell Proliferation.” Proceedings of the National Academy of Sciences of the United States of America 88 (5): 1883–87.

